# An integrated RNA-centric imaging and omics approach reveals distinct properties and composition of neuronal RNA granules

**DOI:** 10.64898/2026.04.27.721160

**Authors:** Jackson A. Rogow, Ella Doron-Mandel, Ronald Cutler, Leti Núñez, Ricardo Vazquez, Laura Harris, Marko Jovanovic, Robert H. Singer, Sulagna Das

**Affiliations:** Dept. of Neuroscience, Albert Einstein College of Medicine, Bronx, NY; Dept. of Biological Sciences, Columbia University, NY; Dept. of Genetics, Albert Einstein College of Medicine, Bronx, NY; Dept. of Cell Biology, Emory University School of Medicine, Atlanta, GA; Dept. of Human Genetics, Emory University School of Medicine, Atlanta, GA

## Abstract

RNA granules are essential regulators of post-transcriptional gene expression, enabling mRNA transport, localization, and local translation in neurons. The localized transcriptome is diverse; however, how different mRNAs are organized into granules for efficient localization and translation remains unknown. Here, we combine real-time endogenous single RNA imaging with protein and RNA proximity labeling to investigate two distinct endogenous neuronal mRNA granule populations, *Actb* and *Arc*, in stimulated primary hippocampal neurons. Using orthogonal RNA labeling systems in a dual knock-in mouse model, we show that *Actb* and *Arc* mRNAs are packaged into spatially segregated granules with distinct trafficking dynamics, localization kinetics, and responses to synaptic stimulation. *Actb* granules displayed rapid and sustained localization, whereas *Arc* granules showed delayed, transient recruitment, consistent with their respective roles in structural and activity-dependent plasticity. Proximity labeling reveals that these granules are distinct in their mRNA composition, despite sharing core RNA-binding proteins, suggesting that shared cis-regulatory elements within mRNA 3’UTR regions drive selective co-packaging of mRNAs into unique granules. Together, these findings demonstrate that neuronal mRNAs are differentially sorted into molecularly and functionally distinct granules, providing a framework for understanding how precise spatio-temporal control of mRNA localization and translation is achieved across complex neuronal arbors.

## Introduction

Post-transcriptional gene expression is centered around the interactions between RNAs and RNA binding proteins (RBPs)^1,2^. These interactions form molecular assemblies known as RNA granules, which are membrane-less structures composed of various regulatory factors depending on the type of RNA granule and the life cycle of the RNA^3–5^. Among these, transport granules are particularly prevalent in highly polarized cells like neurons, where they enable mRNAs to travel relatively long distances and localize to distal dendritic and axonal compartments before being translated at precise locations and times^6^. By delivering mRNAs to remote sites, RNA granules facilitate on-demand protein synthesis distal to the soma, enabling neurons with complex arborizations to adjust their local proteomes rapidly and efficiently^7^.

Tight regulation of these neuronal RNA granules is critical for neuronal development and synaptic plasticity. Altered granule composition and properties results in dysregulated local translation and are associated with a myriad of neurodevelopmental and neurodegenerative disorders, including Fragile X Syndrome and Alzheimer’s disease^7–12^. Hence, understanding the granule composition in healthy and diseased states is crucial to understanding the pathways regulating post-transcriptional gene expression in neurons.

Many neuronal mRNAs are localized in the neuropil as granules^6,8,13–15^, however it remains unknown how these different mRNAs are packaged into granules for efficient transport and localization. Specifically, are the granules homotypic or heterotypic in nature (i.e. contain many copies of the same transcript as opposed to a population of different transcripts), and what kinds of mRNAs are assembled and co-regulated by their respective RBPs within distinct granule populations. Imaging approaches have identified several species of mRNA transport granules that undergo robust activity-dependent trafficking and local translation in the neural arbors in response to stimulation^16,17^. However, whether these behaviors differ between granules and which components drive them remains unclear. Previous studies using biochemical pulldowns have identified components of neuronal RNA transport granules including cognate RBPs, ribosomes, translation factors, motor adaptors, and other regulatory components^6,18^. However, whether granules are generic structures or specific to certain transcripts is unknown.

To date, a systematic comparative investigation of different endogenous RNA granules in living neurons is lacking. Our study compares the dynamics and composition of two endogenous neuronal RNA transport granules and identifies candidates that may drive their behavior. Using an integrated approach that combines live single-molecule imaging with proximity labeling of both RNA and protein in stimulated primary hippocampal neurons, we examined granules containing *Actb* and *Arc* mRNAs — two transcripts that undergo robust activity-dependent transport and local translation and play key roles in synaptic plasticity^16,17,19–22^. To enable simultaneous visualization and biochemical profiling of these endogenous RNAs, we took advantage of the orthogonal MS2 binding sites/MS2 coat protein (MBS/MCP) and PP7 binding sites/PP7 coat protein (PP7/PCP) labeling systems to concentrate fluorescent proteins or the peroxidase APEX2 on the *Actb* and *Arc* transcripts. The MBS/MCP and PBS/PCP systems are orthogonal and exhibit minimal cross-talk and high-affinity binding to their cognate RNA structures^23–26^. We generated a new mouse line whereby both *Arc* and *Actb* mRNAs can be detected in the same neuron using the stem loops knocked into the endogenous 3’UTRs of *Actb* and *Arc* loci^27,28^. Live imaging revealed that *Actb* and *Arc* mRNAs belong to distinct transport granules, exhibit different transport and localization dynamics in response to synaptic stimulation and remain spatially segregated. Proximity-based profiling of these granules showed that they are distinct in their respective RNA and protein interactomes, while sharing many proteins including RBPs likely required for core granule assembly. We identify potential novel candidates for *Actb* and *Arc* granule regulation, including many RBPs and factors related to calcium signaling, translation and degradation. This integrated approach to granule characterization provides insights into the mechanisms by which RNAs are sorted, assembled and licensed for local translation at synapses.

## Results

### *Arc* and *Actb* RNA granules are transported independently along dendrites

A long-standing question has been how neurons package different mRNAs into granules for effective transport and localization. One hypothesis is that functionally related RNAs may be packaged together in the same granule for co-regulation and co-transport. To address this question, we focused on *Arc* and *Actb* mRNAs – two dendritically localized transcripts with established roles in synaptic plasticity^16,17,19–22^.

To interrogate the dynamics of these granules in living neurons, we generated a double knock-in (KI) mouse, where endogenous *Arc* and *Actb* RNAs are labeled with orthogonal stem-loop motifs, allowing for their simultaneous yet distinct detection. Briefly, *Arc*-PBS mice that contain *Arc* mRNAs with 24x PP7 stem loops^27^ were crossed with *Actb*-MBS mice where the *Actb* mRNAs are tagged with 24x MS2 stem loops^28^, to generate the double KI mouse (*Actb*:24xMBS|*Arc*:24xPBS). Each stem-loop motif can be bound by its cognate coat proteins, MS2 coat protein (MCP) and PP7 coat protein (PCP) that are tagged with spectrally distinguishable fluorophores, enabling simultaneous detection of both RNAs in the same cell (Fig. 1A). The 3’UTR tagging does not affect the behavior or regulation of the mRNAs, as determined by previous characterization of individual mouse lines with KI stem loops^27,28^.

**Fig 1.**
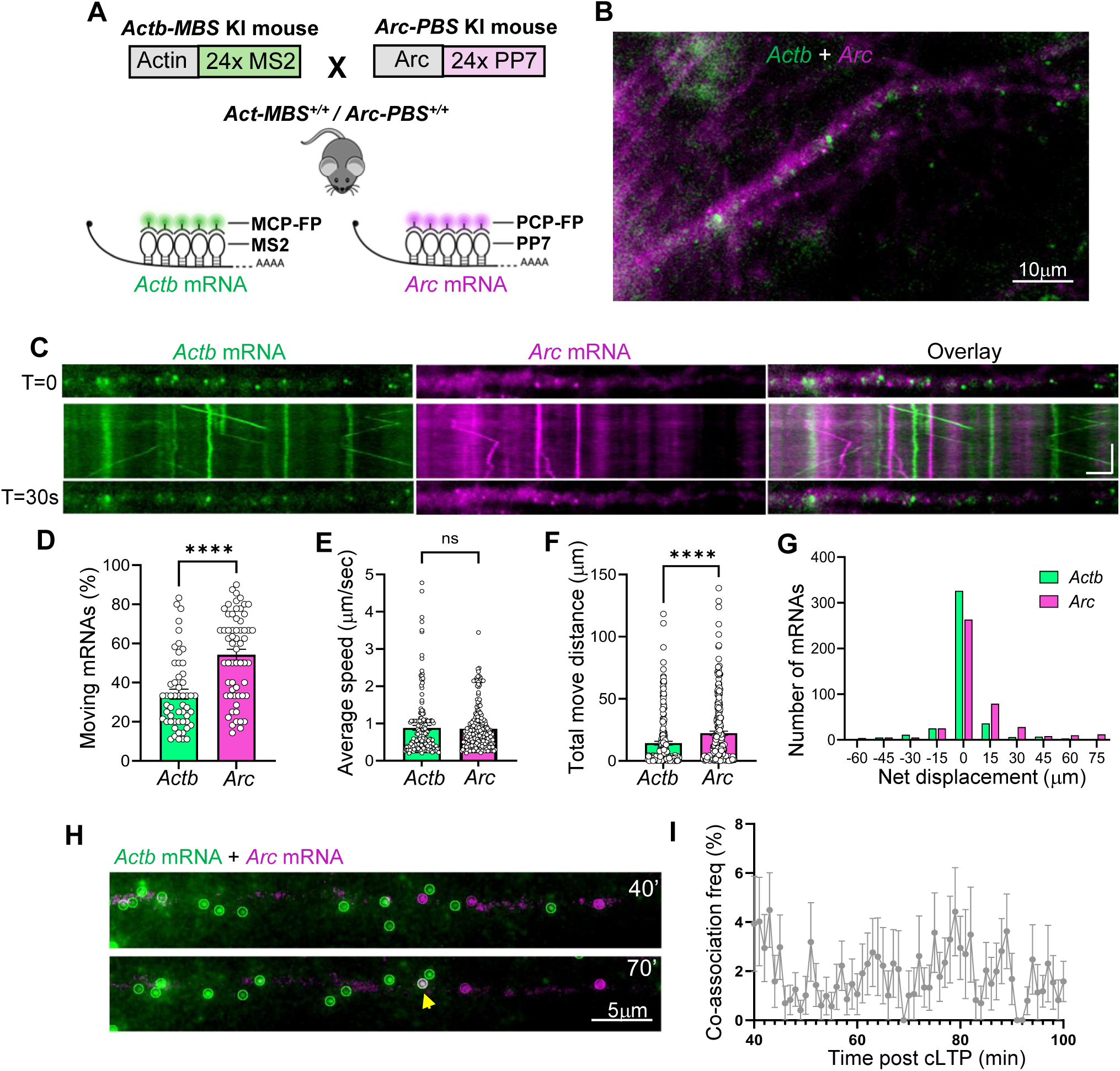
Simultaneous detection of *Arc* and *Actb* mRNAs in individual neurons reveals they are transported independently along dendrites. **(A)** Generation of a double homozygous mouse *Actb*-MBS X *Arc*-PBS that allows detection of both mRNAs in the same mouse using orthogonal stem loops. **(B)** Image of a dendrite from *Actb*-MBS X *Arc*-PBS homozygous mouse showing both mRNAs, *Actb* (green) and *Arc* (magenta). **(C)** Kymographs of *Actb* and *Arc* mRNAs in the same dendrite show distinct tracks. Scale bar= 5microns, 10s **(D)** Fraction of mRNAs moving during the imaging duration (n = 54 dendrites (*Actb*), n= 60 dendrites (*Arc*), p=0.001, Mann Whitney test). **(E)** Average speed of mRNA movement (n = 199 (*Actb*), n= 247 (*Arc*), p=0.11, Mann Whitney test). **(F)** Total distance traversed by the mRNAs during the imaging duration (n = 199 (*Actb*), n= 247 (*Arc*), p< 0.001, Mann Whitney test). **(G)** Net displacement of the mRNAs show anterograde (positive values) versus retrograde (negative values) (n = 421 (*Actb*), n= 442 (*Arc*)). **(H)** Time lapse imaging of *Arc* and *Actb* mRNAs after cLTP (time points indicated), show distinct granules. Colocalization indicated by yellow arrow. **(I)** Frequency of co-localization between *Arc* and *Actb* granules at different time points after cLTP. (n= 11 dendrites, 2 independent experiments). Error bars indicate SEM. ****p<0.001. ns, P > 0.05 is insignificant.

Using this system, we explored the transport and localization behaviors of both *Arc* and *Actb* RNAs in the same neuron. Primary hippocampal neurons cultured from the double KI mouse were transduced with MCP-Halo and PCP-GFP for detection of *Actb*-MS2 and *Arc*-PP7 mRNAs, respectively. Neurons were stimulated by chemical long-term potentiation (cLTP) and imaged 45 minutes (min) post stimulation. Our results showed that both mRNAs localized in distinctly resolved puncta within dendrites (Fig. 1B). Real time imaging of the moving *Arc* and *Actb* granules revealed the two mRNAs were transported entirely independent from each other, often moving on different tracks. (Fig. 1C; Suppl Movie 1). Moreover, these two RNAs had different transport kinetics. While *Arc* granules represented a significantly larger fraction of actively transported mRNAs than *Actb* (Fig. 1D), their speeds were comparable (Fig. 1E). Of note, compared to *Actb*, the total movement of *Arc* mRNAs was higher with a net displacement bias toward the anterograde direction (Fig. 1F,G), suggesting that *Arc* mRNAs may exhibit more directed movement to reach the distal dendrites effectively.

To determine whether these granules retain their distinct identities across time, we performed time-lapse imaging until 100 min post cLTP (Suppl Movie 2). Indeed, most of the mRNAs remained in unique granules, with minimal overlap and co-localization frequencies below 5% across time (Fig. 1H-I). To further assess the co-localization frequency of *Arc* and *Actb* granules across the dendritic length, we performed single molecule fluorescence in situ hybridization (smFISH) using probes against *Arc* and *Actb* (Suppl Fig. 1A). As expected, *Actb* mRNA was significantly more abundant than *Arc* (Suppl Fig. 1B). Notably, the colocalization frequency even in the high density proximal dendrites was ∼3% (Suppl Fig. 1C), which is lower than computed chance interactions between granules and as reported in earlier studies^29^. Together, the findings demonstrate that *Arc* and *Actb* mRNAs form distinct granules that are transported independently of each other, and differ in their transport kinetics and speed. Such spatio-temporal segregation of the granules suggests that they are subjected to distinct post-transcriptional regulation.

### *Arc* and *Actb* mRNAs localize to stimulated spines with unique kinetics

Our observations establish that *Arc* and *Actb* are localized in dendrites in an independent manner. Next, we determined whether these granules respond distinctly to postsynaptic stimulation of the spines. To study the activity-dependent localization of these RNAs, we employed photolytic uncaging of the neurotransmitter glutamate, based on the low-frequency stimulation paradigm^16^, which activates a small subset of spines. Uncaging was done in regions devoid of either *Arc* or *Actb* mRNA to observe the activity-dependent recruitment of mRNAs (Fig. 2A). To obtain robust localization efficiency, uncaging was confined to a 6-micrometers segment with six spots on either side of the dendrite^16^. Both *Actb* and *Arc* mRNAs localized to the stimulated region efficiently, with *Actb* mRNAs exhibiting a higher efficiency of localization compared to *Arc* (Fig. 2A-B). Interestingly, in most cases, *Actb* mRNAs arrived and localized to the uncaged region prior to *Arc* mRNAs and persisted longer (Fig. 2C-D). *Arc* mRNAs transiently localized for 5 min, likely because they were subjected to degradation^30^. Of note, even though *Arc* mRNAs were present in the vicinity of the stimulated region, they did not localize immediately following stimulation (Fig. 2A), suggesting that a precise timing of localization for the granules exists. Interestingly, our results show that even when both *Arc* and *Actb* mRNAs are localized to the same uncaged dendritic segment, their respective granules do not undergo any mixing and retain their distinct properties (Fig. 2E). The unique response of each granule to synaptic stimulation with distinct kinetics of mRNA localization and persistence suggests that *Arc* and *Actb* are independently regulated, likely due to differences in their granule composition and properties.

**Fig. 2.**
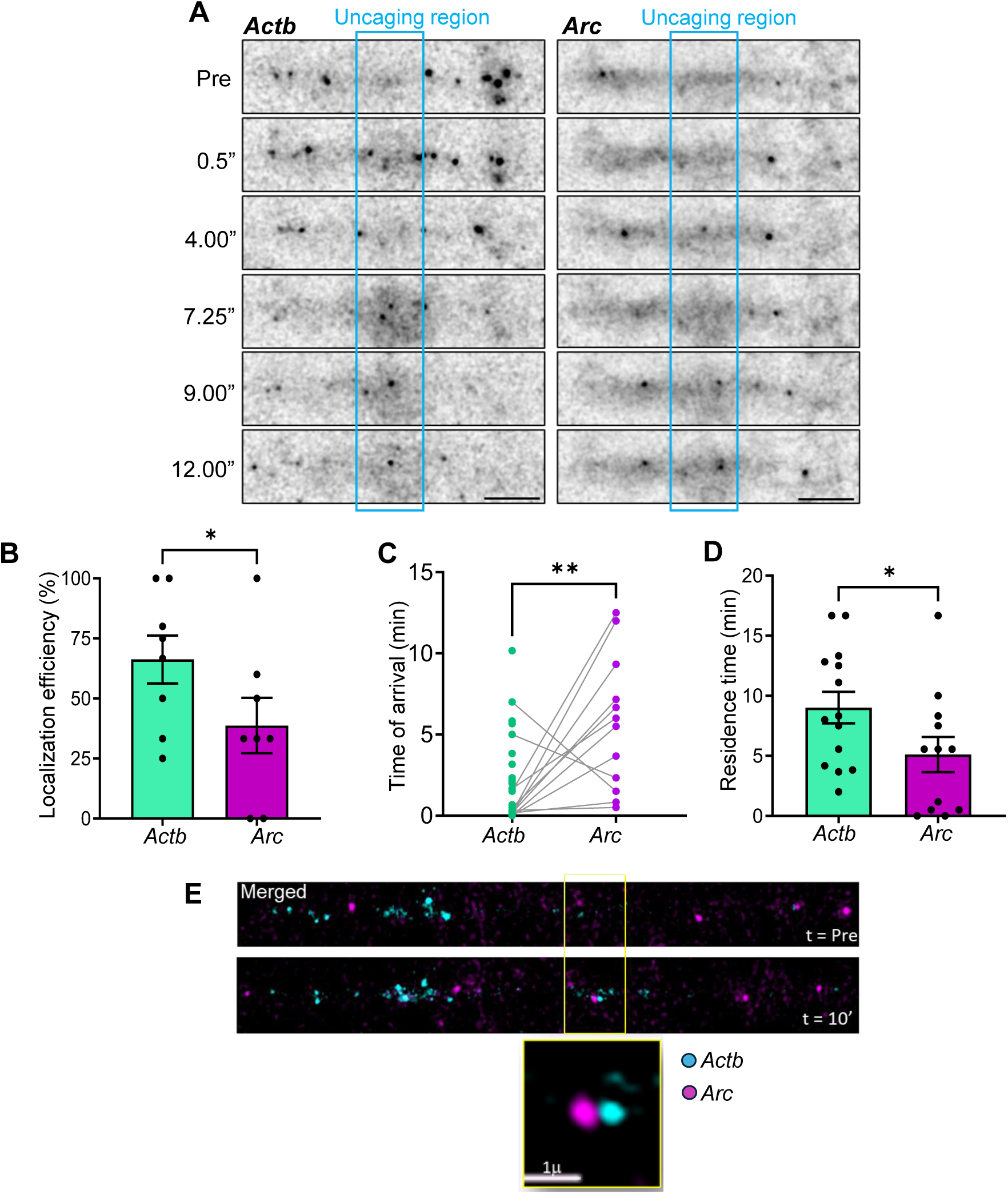
Localization kinetics of *Arc* and *Actb* mRNAs to stimulated spines. **(A)** Time lapse images of *Arc* and *Actb* mRNAs from the same dendrite pre and post uncaging stimulation. Time points after uncaging shown in minutes. Blue outline shows uncaging region. Scale bar 5 μm. **(B)** Localization efficiency of *Arc* and *Actb* in (n = 8 independent experiments, p = 0.046, paired t-test). **(C)** Time of arrival for the mRNA to the uncaged region. Connecting lines show the dendrites where both mRNAs localized (statistical test done only for matched pairs; n = 13 mRNA pairs, p=0.007, Wilcoxon matched-pairs test). **(D)** Residence time of mRNAs at uncaged region (n= 14 dendrites for *Actb*, n=12 dendrites for *Arc*, paired t-test). **(E)** Localization of both *Arc* and *Actb* mRNAs to same uncaging region (yellow outline) but retaining distinct identity. Error bars indicate SEM. *p<0.05. **p < 0.01.

### *Arc* and *Actb* transcripts are packaged with distinct sets of RNAs

The composition of neuronal RNA granules, particularly, whether their RNA composition is homotypic or heterotypic in nature has been debated for many years^6,8^. Our imaging results indicate that, at least for *Actb* and *Arc* mRNAs, each transcript is transported and localized to dendrites within distinct granules. Therefore, we sought to identify the molecular composition of these granules and discover which RNAs, if any, are co-packaged with *Arc* or *Actb* containing granules. To do so, we implemented ascorbate peroxidase (APEX) catalyzed RNA proximity biotinylation followed by sequencing, which is an established method to identify RNAs within a spatial proximity of <20 nanometers to a target inside of a cell^31–33^. Specifically, APEX-based proximity labeling enables rapid, temporally-controlled labeling, facilitating the identification of transient protein and RNA interactions^31,32^. APEX2 was targeted to *Actb* and *Arc* mRNAs using the MBS/MCP and PBS/PCP systems as was used for imaging, with the APEX2 enzyme fused to the PCP or MCP proteins instead of fluorophores^34^. Validation of the APEX2-mediated biotinylation was conducted in wild-type (WT) or endogenously tagged *actb*:24xMBS mouse embryonic fibroblast cell lines expressing integrated coat protein-APEX2 lentiviruses, which showed pulldown of biotinylated RNAs when MCP-APEX2 binds to the 24xMBS and increased total biotinylated protein under reactive conditions (Suppl Fig. 2A-B).

We next implemented this proximity-based labeling in primary hippocampal cultures following cLTP stimulation, matching the conditions used in our imaging experiments. Hippocampal neurons from *actb*:24xMBS and *arc*:24xPBS mice were transduced with MCP-APEX2 or PCP-APEX2 lentivirus, and proximity biotinylation was performed with a 1 min H_2_O_2_ treatment one hour post cLTP stimulation (Fig. 3A). We chose this time post stimulation to capture the maximum number of cytoplasmic granules since it coincides with the peak *Arc* mRNA accumulation in the dendrites, and lowest nuclear/cytoplasmic ratio^17,27^. To account for non-specific biotinylation and labeling, besides the uninfected negative controls, we included “scramble” controls whereby MCP-APEX2 is expressed in *Arc*:24xPBS neurons and PCP-APEX2 is expressed in *Actb*:24xMBS. Following labeling, total RNA was isolated, biotinylated RNAs were enriched by streptavidin bead pull down and libraries were prepared for sequencing (Fig. 3A, Methods).

**Fig 3.**
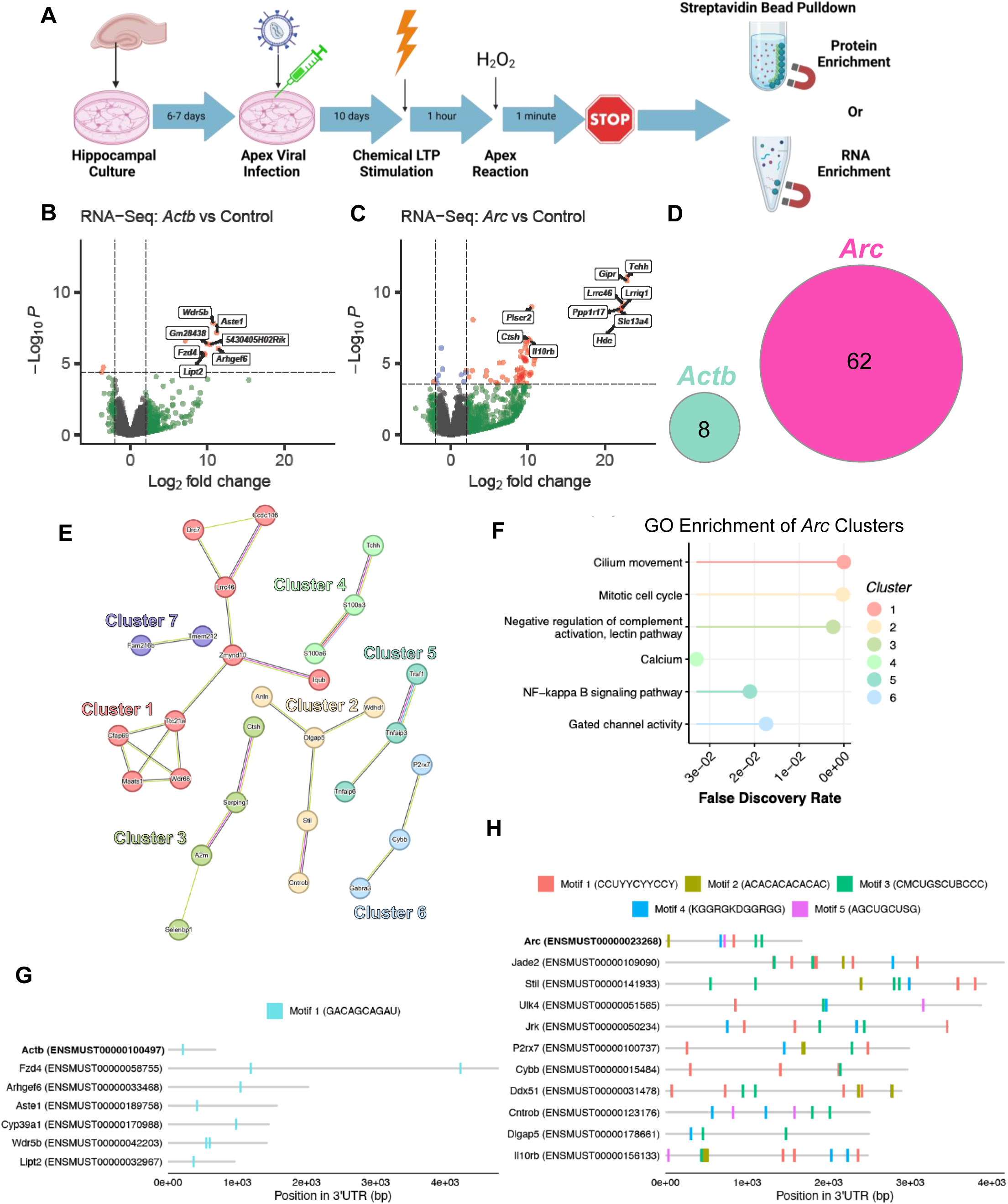
RNA proximity biotinylation shows *Arc* and *Actb* transcripts are packaged with distinct sets of RNAs. **(A)** Experimental pipeline, created with biorender.com^113^ **(B-C)** Volcano plots of **(B)** *actb*:24xMBS proximity labeled RNAs vs Control (no infection + “scramble” PCP-APEX2 virus) and **(C)** *arc*:24xPBS proximity labeled RNAs vs Control (no infection + “scramble” MCP-APEX2 virus). Red points are Log_2_ fold change>2, adjusted p-value<0.05. Statistics: Wald test followed by Benjamini-Hochberg correction for multiple hypothesis testing. **(D)** Venn Diagram showing no overlap of the transcripts enriched in *Arc* and *Actb* granules (Log_2_ fold change>2, adjusted p-value<0.05). **(E)** STRING interaction network of the *Arc* granule enriched transcripts identified in **(C)**. Clusters based on interactions between genes. Interaction lines: Purple = experimentally determined, green = text mining, black = co-expression, blue = curated database, red = gene fusion. **(F)** Over-representation analysis for each cluster as shown in (E) using GO and KEGG databases. Most significant term based on false discovery rate (FDR) is shown for each cluster. Statistics: FDR values were computed using a hypergeometric test followed by Benjamini–Hochberg false-discovery-rate correction. **(G, H)** *De novo* motif enrichment results for *Actb* (**G**) and *Arc* (**H**) enriched transcripts, IUPAC nomenclature. Motifs are shown with matches to 3’ UTR region with q-value<0.1. Statistics: p-values are derived from the probability of equal or higher scoring matches under a background model, with multiple testing correction performed using Benjamini–Hochberg FDR. See Suppl. Fig. 5 for full results.

To identify the RNAs in proximity to *Arc* and *Actb* in each neuronal granule, we performed differential expression analysis comparing each of the enriched samples (i.e. *Actb/Arc*) to their corresponding negative controls (i.e. scramble and uninfected), while filtering out nonspecific hits arising from pull down or viral infection (Suppl Fig. 3, Methods). This analysis identified 8 genes significantly enriched (Log_2_ fold change > 2; padj < 0.05) in *Actb* granules, and 69 genes enriched in *Arc* granules (Fig. 3B-C). Strikingly, the enriched RNAs were unique to each granule, indicating that *Arc* and *Actb* are packaged in distinct RNA transport granules with unique mRNA composition (Fig. 3D).

To determine whether the RNAs packaged in each granule are functionally related, we performed over-representation analysis on the significantly enriched transcripts using the gene ontology (GO) database. While there were no significant GO enrichments for *Actb* (due to the small number of significantly enriched genes), the most significant GO enrichments for *Arc* were cilium movement (Fig. 3E; cluster 1) and mitotic cell cycle (Fig. 3E; cluster 2), containing microtubule regulators and centrosome-associated genes (Fig. 3E-F). Although these annotations were unexpected considering the canonical functions of these genes, most of them encode for microtubule associated proteins, consistent with prior observations that Arc protein participates in cytoskeletal remodeling during structural plasticity and co-sediments with microtubular proteins^21,35,36^.

In addition, several plasticity-related GO terms were observed in the *Arc* enriched transcript clusters including calcium signaling, gated channel activity and inflammatory signaling (complement and NFKb), (Fig. 3E-F, Suppl Fig. 4B-C). *Arc*-specific interactors also included transcripts involved in neurite and dendrite outgrowth (*anln, plcd3, cdc42ep3, tnfaip3, jade2, tppp3, il10rb, selenbp1, ulk4,* and *stil*) and synaptic plasticity (*p2rx7, hdc, gipr* and *gper1*)^37–50^. Like Arc, the G-protein coupled estrogen receptor (*gper1*) regulates AMPA receptor trafficking, and the gastric inhibitory polypeptide receptor (*gipr*) is implicated in learning and memory and is highly upregulated by Arc overexpression^49–51^. These findings suggest that *Arc* mRNA may associate with transcripts sharing common cytoskeletal and plasticity-related functions.

*Actb*-specific interactors included *fzd4* and *αpix/arhgef6*, both involved in cytoskeletal remodeling and dendritic morphogenesis through modulation of Wnt signaling and Rac activity, respectively^52,53^. Given that both *Actb* and *Arc* enriched transcripts contain genes with functions in dendrite structure and plasticity, it is plausible that shared functional roles are not the only criteria for packaging mRNAs together.

Next, we interrogated additional governing principles, other than shared functional roles, that can affect granule assembly. RNAs are often recognized and regulated by RBPs through sequence or structural motifs located in their UTRs^6^. We hypothesized that RNAs that are packaged together in a granule may contain common regulatory motifs. To test this and identify shared cis-regulatory elements associated with each granule, we performed *de novo* motif analysis on the 3’UTR regions of transcripts enriched in either *Actb* or *Arc* granules. One *de novo* motif was identified with the *Actb* associated mRNAs, and strikingly, all mRNAs in this group contained at least one instance of this predicted motif in their 3’UTRs (Fig. 3G, Suppl Fig. 5A). Similarly, 52 out of 61 mRNAs enriched in *Arc* granules contained at least one of the five *de novo* motifs identified in the *Arc* 3’UTR (Fig. 3H, Suppl Fig. 5B, C), suggesting that shared cis-regulatory sequences may contribute to granule-specific recruitment. Importantly, these motifs were largely mutually exclusive between the two transcript groups: none of the 3’UTRs of *Arc* or *Arc*-associated transcripts contained the *Actb* motif, and only one *Actb*-associated transcript contained a single *Arc* motif, further supporting the notion that a cis-mode of regulation may govern co-packaging (Suppl Table 1). We next asked whether the identified *de novo* motifs matched known eukaryotic RBP motifs. Searching against the CisBP-RNA 2.0 database^54^, the *de novo Actb* motif had a significant match for the RBP SRSF4 (Suppl Fig. 5D). Among the *de novo Arc* motifs, significant matches were found for motif 2 and the RBPs HNRNPL, HNRNPLL, RBPMS, RBPMS2, and CNOT4 (Suppl Fig. 5E).

Overall, the RNA proximity biotinylation experiments reinforce our imaging findings, supporting the notion that *Actb* and *Arc* are packaged into molecularly distinct granules containing different repertoires of RNAs. Notably, subsets of transcripts within each granule shared common motifs, suggesting that cis-regulatory elements within these RNAs may guide their selective co-packaging and organization into specific granules.

### *Arc* and *Actb* RNA granules have common and distinct protein components

We next asked which trans-acting protein regulators contribute to the organization of the granules. The protein composition of RNA transport granules remains largely unknown and fundamental questions about how their molecular architecture supports granule-specific behavior are still unresolved. We hypothesized that the repertoire of RBPs regulating each granule type may vary. However, are there also common or shared RBPs between these distinct RNA granules? To test these possibilities, we interrogated the protein composition of each granule. Using the same strategy for profiling RNAs in the granules, we conducted APEX2 mediated proximity biotinylation by directing the APEX2 enzyme to either *Arc*:24xPBS or *Actb*:24xMBS transcripts via fusion to PCP or MCP, respectively. Biotinylated proteins were enriched by streptavidin bead pull down and processed for liquid chromatography tandem mass spectrometry (LC-MS/MS, Methods). Because our primary objective was a direct comparison between *Actb* and *Arc* interactors, these experiments were performed in neurons from the double KI mouse, which express both *arc*:24xPBS and *actb*:24xMBS. Neurons were transduced with either the APEX2-PCP or APEX2-MCP lentivirus, and uninfected cells were used as negative controls to estimate background binding.

Overall, our dataset identified 1992 proteins, including APEX2, which were reproducibly detected in all experimental samples, but not in the negative controls (Suppl Fig. 6). We first identified the protein interactors for each transcript independently by comparing enrichment in each experimental sample (*Actb* or *Arc*) relative to the proteins pulled down from uninfected neurons. We identified 204 *Arc* interactors and 133 *Actb* interactors, including APEX2 itself in both samples (Log_2_ fold change > 2; padj < 0.1, Fig. 4A-B). Expectedly, 56% of all hits and 64% of common hits (125/222 total and 74/115 common) have established roles as RNA regulators, based on comparing our hits against the RBP2GO database^55^. In addition, overrepresentation analysis of the union of *Arc* and *Actb* interactors (222 proteins) using the Gene Ontology (GO) database revealed enrichment of mRNA metabolism terms (Fig. 4C). A similar enrichment was observed for each of the interactomes (*Arc* and *Actb)* independently (Suppl Fig. 7A-B).

**Fig 4.**
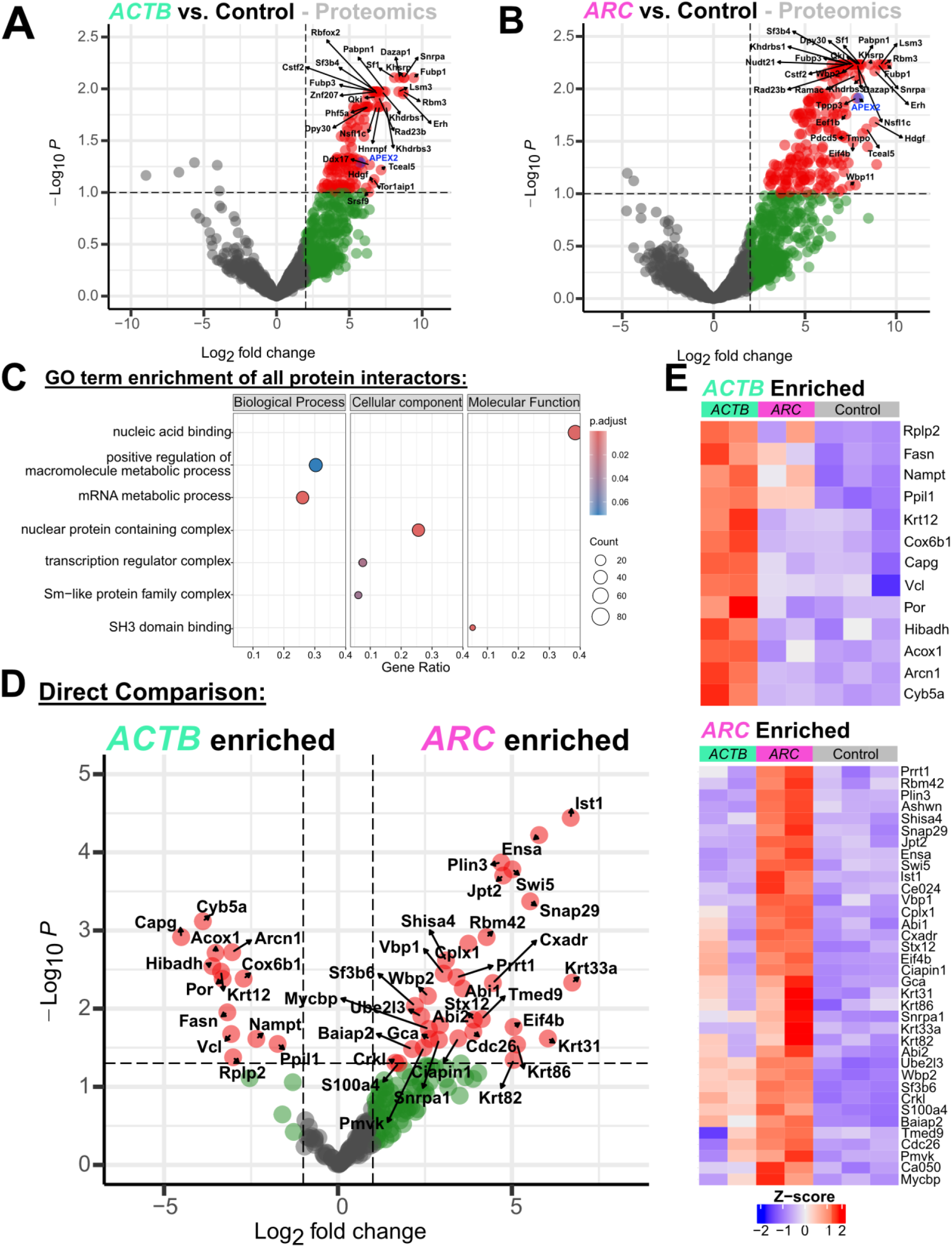
Protein proximity biotinylation identifies shared and distinct protein components of *Arc* and *Actb* granules. **(A-B)** Volcano plots showing the differential analysis between *Actb/Arc* and negative controls to define interactors (Limma statistical model with p-value correction using the Benjamini-Hochberg method). Adjusted p-values < 0.1 and log_2_(Fold Change) > 2 were used as cutoffs to define interactors). Overall, the union of interactors between both *Actb* and *Arc* resulted in 222 proteins (See Suppl Fig. 7). **(C)** GO term enrichment based on the 222 proteins identified as *Actb* or *Arc* interactors. **(D)** Volcano plot showing the direct comparison between *Arc* and *Actb* signals for the 222 proteins found as interactors (Limma statistical model, p-value < 0.05 and log2 Fold Change > 1 were used as cutoffs to define “enriched” interactors). Overall, we identified 36 / 13 enriched *Arc* / *Actb* interactors, respectively. **(E)** Heatmap representation of the signal of enriched *Actb* and *Arc* interactors (as defined in the direct comparison shown in **(D)** across all samples).

We next examined differences in the protein composition of *Arc* and *Actb* granules. To this end, we considered the 222 proteins identified as interactors in either sample (Suppl Fig. 7C) and performed a direct comparison of their signal in *Arc-* versus *Actb-*enriched samples. Because both samples were derived from the same genetic background and cell population, they effectively serve as reciprocal controls for one another. Using a p-value cutoff of <0.05 and a log_2_fold-change >1, we identified 36 *Arc*-enriched and 13 *Actb*-enriched proteins that were selectively associated with each granule type (Fig. 4D-E).

Among the 173 hits that were not significantly different between *Arc* and *Actb* (Suppl Fig. 7D), we detected proteins previously reported to regulate both *Arc* and *Actb* mRNAs. For example, Sam68 (khdrbs1), an RBP implicated in synaptic plasticity and dendritic regulation of both transcripts^56,57^ as well as its interaction partners, Khdrbs2 and Khdrbs3. As previously mentioned, the common protein interactors also included many RBPs, including six members of the Heterogeneous nuclear ribonucleoprotein (Hnrnp) family, five members of the small nuclear ribonucleoprotein family, and others (Detailed lists of the proteins in each group are provided in Suppl. table 2). In addition, we found multiple translation regulators such as Eif4h, Eif2s3x, Eef1b, Rbm3, together with several proteasome components. Interestingly, several calcium-binding proteins were also present among the shared interactors of both RNAs, including Calm1/2/3, Hpcal4, Pef1 and its interactor Pcdc6, Vsnl1, and S100a16, many of which participate in neuronal calcium signaling^58^. These findings support the notion that RNA transport granules contain calcium sensing machinery enabling them to sense and respond to local changes in intracellular calcium associated with synaptic activity.

Next, we examined the functional relationships within each group of interactors using the STRING database^59^ (Suppl Fig. 7E-F). This analysis revealed two clusters of metabolic enzymes (Fasn, Acox1, and Hibadh; Cyb5a and Por) among the *Actb* enriched interactors. In contrast, *Arc* enriched interactors formed several functional clusters by STRING, including one group of vesicle associated SNARE family proteins (Stx12, Snap29, Cplx1) and a cluster involved in cytoskeletal remodeling (Abi1, Abi2, and Baiap2/IRSp53). Our data showing Baiap2 interacts with *Arc* mRNA is in agreement with recent evidence showing that Baiap2 co-localizes with Arc in response to stimulation, and is important for biogenesis and release of *Arc* mRNA containing extracellular vesicles^60^.

Together, these results indicate that *Arc* and *Actb* granules share a substantial core set of proteins, many of which function in RNA metabolism and translational control. Alongside this, the identification of granule-specific protein components supports a model in which a subset of specialized interactors may contribute to distinct stimulus responsiveness and spatio-temporal dynamics of *Arc* and *Actb* granules.

## Discussion

Localization and local translation of mRNAs plays a central role in shaping the local proteome, enabling neurons to efficiently form and remodel specific connections for synaptic function and plasticity^61–63^. With a growing catalog of dendritically localized mRNAs and their RBPs, a combinatorial challenge is presented in how these mRNAs are selectively packaged into transport granules for trafficking, localization and translational control. In particular, the molecular logic governing RNA granule assembly, transcript co-packaging, and activity-dependent remodeling of these assemblies has remained unresolved because of technical limitations to profile the endogenous transport granules. Here, using an integrated strategy that combines live single RNA imaging with proximity-based transcriptomics and proteomics, we defined the real-time dynamics and molecular architecture of endogenous RNA transport granules. Focusing on *Arc* and *Actb*, two plasticity-relevant transcripts that are localized to neuronal dendrites, we demonstrate that these granules are fundamentally distinct in their transport dynamics, localization kinetics, responsiveness to stimulus, and molecular composition.

Simultaneously imaging both transcripts within the same neuron revealed that the *Arc* and *Actb* granules are spatially segregated, and localize independently in response to both local and global synaptic stimulation. Compared to *Actb* granules, *Arc* were more dynamic and exhibited greater anterograde displacement but were recruited slowly to the stimulated spines. These properties may be related to Arc’s role as an immediate early gene (IEG) with short mRNA half-life (∼30-60min), and its ability to undergo recurrent localization to dendritic translation hubs to confer specificity during spine remodeling^17,64–66^. In contrast, dendritic *Actb* mRNA persisted at stimulated spines, likely supporting multiple rounds of translation^16,67^. Although both mRNAs are recruited to stimulated spines following glutamate uncaging, their distinct temporal localization patterns indicate that RNA granules respond differently to local signaling, likely due to differences in their molecular composition.

Previous work has suggested that neuronal RNA granules may contain mixed RNA species assembled through shared RBP binding, whereas others studies indicate independent RNA localization^68,69^. To resolve this question in an unbiased manner, we employed RNA-targeted proximity-based transcriptomics and found that *Arc* and *Actb* mRNAs associate with non-overlapping RNA repertoires. At the same time, we find evidence that subsets of functionally related RNAs interact and enrich within the same granule population, suggesting that shared functional relationships may contribute to RNA co-packaging. This notion is consistent with a recent large-scale granule profiling study that found protein coding RNAs can be co-organized via common function^70^. Notably, *Arc* granule enriched RNAs clustered into plasticity-related pathways, including calcium signaling, inflammatory signaling, and ligand-gated channel activity, as well as two clusters associated with cilia, centrosomes and microtubule regulation. Although the latter categories were unexpected, emerging evidence indicates that the centrosome can act as an RNA regulatory hub in mature neurons and centrosome and ciliary pathways are involved in memory formation in the hippocampus^71,72^. These findings suggest that *Arc* granules can coordinate cytoskeletal and signaling programs relevant to activity-dependent remodeling. In addition, we identified both known and novel sequence 3’UTR cis-regulatory elements among transcripts associated with either *Arc* or *Actb* granules, providing insights into how different RNAs could be packaged together via shared sequence motifs.

Our proteomic analysis revealed that *Arc* and *Actb* granules share most proteins while also containing distinct sets of regulatory components, with the majority belonging to RBPs. Since RBPs are pleiotropic and frequently operate within combinatorial regulatory networks across sub-cellular compartments^3,8,73–77^, partial overlap between granule proteomes is expected. All mRNAs go through a general metabolic sequence from transcription to degradation involving many canonical proteins, and thus two different mRNA species will interact with common effector proteins across their life cycles^78^. The shared proteome included canonical translation regulators such as Eif4h and Eef1b^79,80^, as well as RBP modulators like Peptidyl-prolyl cis-trans isomerase NIMA-interacting 1 (Pin1)^81^, suggesting that common factors controlling RNA metabolism can form a conserved scaffold for granule organization. In addition, we identified many regulatory factors that may modulate granule properties via different signaling mechanisms. For example, both granules contained multiple calcium binding proteins (such as Calm1/2/3 and Vsnl1), supporting the notion that transport granules may function as activity-responsive signaling modules capable of sensing local calcium changes^58^.

Despite this shared architecture, we identified granule-specific protein components that likely confer functional specialization. *Arc*-enriched proteins were associated with synaptic plasticity and vesicle/membrane trafficking, whereas *Actb*-enriched interactors included metabolic enzymes. Of note, *Arc* enriched protein interactions include the RBPs Rbm42, Eif4b and Sf3b6, which have demonstrated roles in mRNA translation, as well as other synaptic, cytoskeletal, and membrane trafficking proteins^82–84^. *Actb* enriched interactors Fatty acid synthase (Fasn) and calcium dependent actin capping protein Capg^85^ have both been identified as having potential RNA-binding activity in previous large scale omics studies^55,86–89^. These differences suggest that *Actb* granules are more aligned with structural maintenance programs, whereas *Arc* granules preferentially coordinate activity-dependent synaptic signaling pathways.

Notably, the absence of large sets of RBPs uniquely restricted to either granule raises the possibility that granule identity emerges from combinatorial interactions among shared RBPs, and/or additional regulatory layers mediated by the signaling proteins and post-translational modifications. For example, even shared RBPs could drive independent localization and regulation of granules guided by differences in RNA-RBP stoichiometry or binding configurations^90^. Other regulatory mechanisms via post translational modifications, such as phosphorylation, methylation, sumoylation, and ubiquitination, could modify RBPs and RNA-RBP interactions, therefore resulting in distinct granule properties despite partially shared molecular components^4,90–93^. Together, these findings support a model in which RNA granules are assembled through a shared regulatory core that is tuned by transcript-specific protein interactions and activity-dependent signaling to generate functionally distinct transport assemblies.

In conclusion, we establish a framework for understanding how neuronal RNA granules are assembled through RNA-centric regulatory logic and dynamically shaped by neuronal activity. Given that granule dysregulation is implicated in neurological disorders, future directions investigating how perturbation of granule components and disease causing RBP mutations impacts localization, organization and behavior of neuronal granules will be critical^9,10,94–97^. Broadly, our study demonstrates how integrating high-resolution imaging with proximity omics enables the direct dissection of the principles governing RNA granule organization in living neurons and lays the foundation for understanding post-transcriptional control in health and disease states.

## Methods

### Primary Mouse Hippocampal Neuron Cultures

Hippocampi were isolated from *Actb*:24xMBS|*Arc*:24xPBS*, Arc-PBS*, *Actb*-MBS^M/M^ mice on post-natal day 0 (P0), and both sexes were used and combined. Hippocampal tissue was digested with 0.25% trypsin for 15 minutes at 37℃, triturated and plated onto Poly-D-Lysine (Sigma) coated 6 well plates for APEX experiments or glass bottom Mattek dishes for imaging experiments. For APEX experiments cells were plated at a density of 650,000 per well in 6 well plates, and each processed sample for downstream experiments consisted of 2 harvested wells combined of the same litter, genotype and viral conditions. For imaging experiments, cells were plated at 75,000 cells/dish for live imaging and 60,000 cells/dish for fixed cell imaging. After plating, neurons were incubated for 2 hours in Neurobasal A media with 10% FBS (R&D systems), GlutaMax (ThermoFisher), and Primocin (InvivoGen) at 37C. Following this, neurons were grown in Neurobasal A media supplemented with 1% FBS (R&D systems) B-27 (ThermoFisher), GlutaMax (ThermoFisher), and Primocin (InvivoGen).

### Plasmids and viruses

The NLS-HA-stdMCP-3xFLAG-APEX2-P2A-yGFP and NLS-HA-stdPCP-3xFLAG-APEX2-P2A-yGFP plasmids used for all APEX experiments (referred to as MCP-APEX and PCP-APEX respectively) were cloned using standard molecular biology techniques (stdMCP and stdPCP are synonymized tandem dimer coat proteins). For live imaging, plasmids expressing both coat proteins from the same construct were used: NLS-HA-tdMCP-tagRFPt-IRES-NLS-flag-tdPCP-GyG, and NLS-HA-tdMCP-GyG-IRES-NLS-flag-tdPCP-Halo.

Lentiviral particles for hippocampal APEX experiments were generated by CaCl_2_ mediated transfection of expression plasmids along with RRE, REV, and VSVG plasmids into HEK 293T cells, followed by collection with Lenti-X concentrator (Takara). APEX Virus aliquots were tested on small batches of neurons to approximate MOI via yGFP visualization.

### Lentiviral-Generated Stable Cell lines

To generate stable cells lines with the plasmids, lentiviruses were generated by transfecting HEK293T cells with Lipofectamine 2000 (Thermo Fisher) based on manufacturer’s recommendations. Harvested lentivirus from HEK293T cells was filtered with 0.45 µm filter. and collected with Lenti-X concentrator (Takara Bio) using an overnight incubation. Solution was centrifuged at 1500 x g for 45 minutes at 4 degree Celsius and supernatant was resuspended at (1:10 or 1:100) in DMEM with 10% fetal bovine serum and 1% penicillin/streptomycin. Lentiviruses were stored at -80 degrees Celsius prior to use. β-actin MBS homozygous or +/+ immortalized mouse embryonic fibroblasts^28^ were incubated with 10 µm/mL polybrene and infected with 500 µL of MCP-APEX or PCP-APEX lentiviruses and incubated for 48-72 hrs. After incubation, cells were sorted by green fluorescent protein expression using fluorescent activated cell sorting.

### Polymerase Chain Reaction with Gel Electrophoresis

Reverse transcriptase PCR and gel electrophoresis performed on biotinylated RNA enriched by streptavidin bead pulldown as described in the RNA isolation and enrichment section. cDNA was synthesized using Superscript III (Invitrogen). *ACTB* and *GAPDH* primers were used for PCR as described in Eliscovich et al. and Lionnet et al.^28,29^:Actb_MBSFwd:5′-GATCTGCGCGCGATCGATATCAGCGC-3′; Actb_MBSRev:5′-GCCAGCCCTGGCTGCCTCAACACCTC-3′; GAPDHFwd:5′-GAGCGAGACCCCACTAACATCAAATG-3′; GAPDHRev:5′-CAGGATGCATTGCTGACAATCTTGAG-3′.

### Biotinylated Protein Probing

Using the Lentiviral-generated, β-actin MBS, MCP-APEX fibroblast line biotinylated protein was isolated and probed from APEX reactions or no reaction controls using the methods described in Lam et al.^98^ on total protein lysate using Pierce Ripa buffer (Thermo Fisher), and streptavidin-HRP (1:3000, Thermo Fisher).

### stdMCP-3xFLAG-APEX2Immunofluorescence

Cells were plated onto 10 µg/mL fibronectin coated 18-CIR coverslips (Fisherbrand) in 12-well culture plates. Cells were incubated at 37 degrees at 5% CO2 overnight. The next day cells were fixed with 4% paraformaldehyde in 1X Phosphate buffered saline (PBS) for 20 minutes followed by three washes 5 minutes each. After washes, coverslips were quenched using 50 mM glycine (Sigma) in 1X PBS for 5 minutes. Next, coverslips were permeabilized with 0.1% triton x 100 for 15 minutes followed by three washes for 5 minutes each. Samples were blocked with 2% Bovine Serum Albumin in 1X PBS for 30 minutes. After blocking, coverslips were incubated with primary antibody (Mouse Anti-Flag M2 (Sigma, cat#F3165); 1:1000 in blocking solution) for 45 minutes at room temperature. Coverslips were washed with 1X PBS four times 5 minutes each. Secondary antibody was incubated for 40 mins at room temperature in the dark (Life Technologies, Anti-mouse AlexaFluor647; 1:1000 in blocking solution). Washed 3 times with PBS at room temperature, 5 minutes each. Mounted coverslips using mounting medium (25% DAPI in prolong diamond, Life Technologies).

### Chemical Longterm Potentiation Protocol and APEX labeling

Neurons were infected with MCP-APEX2-P2A-yGFP or PCP-APEX2-P2A-yGFP lentiviruses on day 6-7 depending on visual maturity of the culture. Neurons proceeded to have the APEX experiment performed 10 days post infection. On the day of Apex labeling, media was changed to 50µM APV (Tocris) in Hib A with Mg^2+^ (Thermo Fisher) 2 hours before induction. To induce neurons, cultures were incubated with Mg^2+^ free Hib A media supplemented with 200 µM glycine (Sigma) and 100 µM Picrotoxin (Tocris) for 5 minutes. After 5 minutes this media was slowly washed out with 3 washes of Hib A with Mg^2+^. Neurons were incubated for 30 minutes @ 37C and then biotinyl tyramide (Sigma) in Hib A with Mg^2^ was added to each culture to a final concentration of 500 µM biotinyl tyramide. Neurons were incubated for another 30 minutes at 37C and then immediately used in an APEX reaction. To begin the APEX reaction, hydrogen peroxide (Sigma) was spiked into each well to a final concentration of 1mM and cultures were gently swirled on a tabletop rotator for one minute. Media was then quickly decanted and 2X Quench Solution (Sodium Ascorbate 20 mM, Sodium Azide 20mM, Trolox 10 mM, 1x PBS) (Sigma, 10x PBS from Invitrogen) was added to each well and swirled on the rotator for one minute. Cultures were then washed with 1X Quenching Solution (Sodium Ascorbate 10 mM, Sodium Azide 10mM, Trolox 5 mM, 1x PBS) twice for one minute each. Then samples were collected via gentle trituration with Trizol (Invitrogen) and flash frozen in liquid nitrogen for storage at -80C.

### RNA isolation and enrichment with streptavidin beads

RNA was extracted from TRIzol Reagent (Invitrogen) as per the manufacturer’s instructions, using 10µg RNAse free Glycogen (Thermo Scientific) as a carrier. RNA concentration was quantified with nanodrop and samples were treated with Turbo DNAse (ThermoFisher) for 10 minutes at 37℃. DNAse reaction was quenched with TRIzol and samples were purified using the direct-zol RNA miniprep kit (Zymogen) skipping the optional DNase step in the kit. An aliquot of each RNA sample was used to check RNA integrity via a tape station or bioanalyzer (agilent). Only RNA samples with RIN/RQN values above 8.5 were processed for biotinylated RNA enrichment and RNA sequencing. Biotinylated RNA enrichment was performed similar to as described in stress granule APEX-seq^1^ All enrichment buffers were made fresh from RNAse free ingredients whenever possible and all washes occurred on a magnetic rack. Briefly: 10µl of Dynabeads MyOne Streptavidin C1 beads (ThermoFisher) per sample was washed three times with 1mM MgCl_2_, 0.25% sodium deoxycholate, 1x PBS wash buffer. Beads were then washed once and blocked for 30 minutes in 5x Denhardt’s reagent, 150 µg/mL Poly IC, 1mM MgCl_2_, 1x PBS, 0.25% sodium deoxycholate, 1:100 Ribolock RNAse inhibitor blocking buffer. Block was removed and RNA samples were added along with fresh blocking buffer. Samples were incubated on a slow rocking nutator at RT for 1.5hrs ensuring sample mixing. Beads were washed twice with 6M urea, 0.1% SDS, 1x PBS. Beads were washed once with 2% SDS, 1x PBS. Beads were washed once with 750 mM NaCl, 0.25% sodium deoxycholate, 0.1 % SDS, 1x PBS. Beads were washed once with 150mM NaCl, 0.25% sodium deoxycholate, 0.1 % SDS, 1x PBS. RNA was eluted from the beads with 300uL TRIzol and purified with the Direct-Zol RNA direct-zol RNA microprep kit (Zymogen). RNA was not quantified after pulldown, it was frozen at -80C or immediately processed for first strand cDNA synthesis using the Smartseq v4 mRNA Library prep kit (Takara) with maximum RNA input volume. .

### Protein isolation and enrichment with streptavidin beads

Protein was extracted from TRIzol per the manufacturer’s instructions, omitting the protein dialyzing step. After airdrying, the protein pellet was resuspended in RIPA buffer (50mM Tris pH 7.5, 150mM NaCL, 1% NP40, 0.5% Deoxycholate, 0.2% SDS, 1mM DTT, 50x Protease Inhibitor Cocktail), flash frozen, vortexed and bath sonication for 5 minutes. To solubilize the pellet completely, samples were diluted 1:2 to a final 4M Urea concentration with 8M Urea buffer (8M urea, 75mM NaCl, 50mM Tris, 0.5 mM EDTA), bath sonicated for 5 minutes and incubated at 1000 rpm for 1 hr at 50℃. Samples were then spun for 10 minutes at 10,000g 4℃ and the supernatant containing resuspended proteins was transferred to a new tube. Dynabeads MyOne Streptavidin C1 beads (Invitrogen) were used for the enrichment. 10uL beads per sample were first equilibrated with 2x washes of wash buffer (50mM Tris PH 8, 150mM NaCl, 2M urea, 0.2% NP40). Samples were diluted 1:2 with sample dilution buffer (50mM Tris, 150mM NaCl, 50x Protease Inhibitor Cocktail) and added to the beads. Samples rotated overnight at 4℃ with beads. Samples were then washed 4x with wash buffer on a magnetic rack at room temperature, and the proteins were digested off the beads using 80µL of digestion buffer (50mM Tris pH 7.5, 2M Urea, 1mM DTT, 5µg/mL Trypsin) shaking at 1000rpm for 1 hr at 25℃. Samples were placed back on the magnetic rack, supernatant collected, and two final washes with 60uL (2M Urea 50mM Tris pH 7.5) performed. The digest supernatant and supernatant from final washes were combined together, spun to remove trace beads, and moved to a fresh tube. Proteins were then digested overnight at 25 °C and 600 rpm using 0.5 μg of sequencing-grade modified trypsin (Promega). After digestion, peptides were acidified with formic acid (Thermo Scientific) and desalted using in-house packed C18 StageTips (two plugs), following the protocol described by Rappsilber et al^99^. Cleaned peptides were dried using a Thermo Savant SpeedVac and reconstituted in 3% acetonitrile/0.2% formic acid prior to LC-MS/MS analysis.

### LC-MS/MS analysis on a Q-Exactive HF

About 1 μg of total peptides were analyzed on a Waters M-Class UPLC using a IonOpticks Aurora ultimate column (1.7 µm, 75 µm x 15 cm) coupled to a benchtop Thermo Fisher Scientific Orbitrap Q Exactive HF mass spectrometer. Peptides were separated at a 300 nL/min flow rate with a 96-minutes gradient, including sample loading and column equilibration times, using solvents A (0.1% formic acid in water) and B (0.1% formic acid in acetonitrile). The detailed gradients of solvent B are: 2% B for 1 min; linear increase to 10% B over 30 min; linear increase to 22% B over 27 min; linear increase to 30% B over 5 min; linear increase to 60% B over 4 min; linear increase to 90% B over 1 min, held for 2 min; linear decrease to 50% B over 1 min, held for 5 min; linear decrease to 2% B over 1 min; and re-equilibrated at 2% B for the rest of the acquisition period. Data were acquired in data-dependent mode using Xcalibur software (4.5.474.0). MS1 spectra were measured with a resolution of 120,000, an AGC target of 3e6, and a scan range from 300 to 1800 m/z. Up to 12 MS2 spectra per duty cycle were triggered at a resolution of 15,000, an AGC target of 1e5, an isolation window of 1.6 m/z and a normalized collision energy of 25.

### Data Analysis for proteomics

All raw data were analyzed with SpectroMine software version 16.0.220606.53000 (Biognosys) based on a UniProt mouse database (release UP000000589, Mus Musculus, Taxon ID 10090), performed with the “BGS factory settings” including the following parameters: Oxidation of methionine and protein N-terminal acetylation as variable modifications; carbamidomethylation as fixed modification; Trypsin/P as the digestion enzyme; For identification, we applied a maximum FDR of 1% separately on protein and peptide level. “Cross run normalization” and “Global imputing” were inactivated. This gave intensity values for a total of 1,992 protein groups across all samples and replicates. “PG.Label-Free Quant” values were used for all subsequent analyses.

### RNA-sequencing and transcriptome analysis

Libraries were prepared with Takara SMART seq v4 Libraries for mRNA analysis using 1ng cDNA input and 16 cycles for the library amplification step, and sequenced using 150 paired-end mode to ∼20 million reads per sample on an Illumina Novaseq platform by Novogene Corp Inc., CA. Raw sequence reads were processed using the nf-core RNA-Seq pipeline^100,101^ to obtain gene abundance levels.

All analyses were performed at the gene level using the gene abundance values produced by RSEM^102^. Genes were filtered out if the measured transcript per million (TPM) < 1 amongst all experimental replicates. (Supplementary Figure 3A) Correlation analysis of TPM values between replicate samples was performed to ensure high concordance (Supplementary Figure 3B). *Actb* and *Arc* samples were analyzed separately, due to large differences in the amount of genes detected (Supplementary Figure 3C). Gene count normalization and differential gene abundance analysis was performed using DESeq2^103^. For comparisons of *Actb/Arc* vs. control samples, control samples included both the uninfected control sample and the relevant stem loop matched, virus scrambled control sample. Genes were considered to be significantly enriched if the Log_2_ fold-change > 2 and adjusted p-value < 0.05. Technical artifacts were filtered out of the lists of significantly enriched genes based on differential expression analysis on the effect of the pull-down (Supplementary Figure 3D) and effect of the viral infection (Supplementary Figure 3E). Sequences of the MCP and PCP viruses as well as the endogenous locus stem-loop containing sequences for *Actb* and *Arc* were included in the alignment (Supplementary Figures 3F-G).Over-representation analysis was performed by clusterProfiler^104^. Network analysis was performed using the STRING database where genes which had no interactions in the resulting network were filtered out. Sub-networks of genes were identified in the resulting network using K-means clustering. Gene ontology enrichment analysis was performed on each sub-network^59^.

### 3’ untranslated region motif analysis

3′UTR sequences were extracted from the Ensembl mouse annotation (GRCm39.113) and GRCm39 primary genome assembly. For each gene, the longest annotated 3′UTR isoform was retained. The foreground set comprised transcripts identified as enriched in the APEX-seq *Actb* or *Arc* versus uninfected and scramble comparisons (log2fc > 2, padj < 0.05), with *Actb* or *Arc* added manually before extraction. The background set comprised transcripts classified as nonenriched in the corresponding APEX-seq comparison (log2fc < 0, padj > 0.05). *De novo* motif discovery was performed with MEME Suite (version 5.5.8) using STREME^105^, DREME^106^, and MEME^107,108^ in RNA mode, with enriched 3′UTRs used as the foreground set and nonenriched 3′UTRs used as the zero-order background set. Motifs identified across discovery runs were merged and deduplicated and visualized with universalmotif ^109^.

To compare *de novo* motifs with known RNA-binding protein motifs, the deduplicated motif set was queried against the CisBP-RNA 2.0^54^ MEME-format reference database with TomTom^110^. Significant matches between *de novo* and reference motifs were visualized with universalmotif^109^. Motif occurrences across enriched 3′UTRs were mapped with FIMO^111^ using the deduplicated *de novo* motifs where matches with q-value < 0.1 were retained.

### Single molecule FISH and imaging

The detailed protocol for single-molecule FISH has been published before^67^. Briefly, cells were fixed, permeabilized, and incubated overnight at 37 °C with 100 nM probes against MS2 and PP7 stem loops, for MS2: 5’-C**T**GCAGACATGGG**T**GATCCTCATGTTTTC**T**A-3’ and for PP7: 5’-C**T**GCAGGGAGCGACGCCA**T**ATCGTCTGCTC CTT**T**C-3’ (bolded Ts indicate modifications for dye conjugation). MS2 probes were labeled with Cy5 and PP7 probes with Cy3. Following hybridization of probes and washes, samples were mounted using ProLong Diamond antifade reagent containing DAPI (Life Technologies).

Images were acquired using a custom upright wide-field Olympus BX-63 microscope equipped with an X-Cite 120 PC illumination system (EXFO), an ORCA-R2 CCD camera (Hamamatsu), and a SuperApochromatic 60×/1.35 NA objective (UPLSAPO60XO). Zero-pixel-shift filter sets (Semrock; DAPI-5060C-Zero, Cy3-4040C-Zero, Cy5-4040C-Zero) were used for multichannel imaging. Images were collected at a pixel size of 107.5 nm in XY with 200 nm Z-steps.

### Live Cell Imaging

Hippocampal neurons were transduced with viruses expressing stdMCP-Halo-IRES-stdPCP-GyG, or with stdMCPtagRFPt-IRES-stdPCPGyG. Mattek dishes were switched to Hibernate A media the night before imaging, and for cLTP experiments APV (50μM) was added overnight. Halo dyes (JF646) was added at 10 μM for 30 min, followed by washes and treatment for cLTP.

Time-lapse imaging of cultured mouse hippocampal neurons was performed using a wide-field fluorescence microscope built on an Olympus IX-81 platform equipped with an iXon Ultra DU-897U EMCCD camera (Andor Technology Ltd., Belfast, UK) and a UPlanApo TIRF 150×/1.45 NA oil-immersion objective (Olympus). Excitation was provided by 491-nm (Calypso-25, Cobolt), 561-nm (LASOS-561-50), and 640-nm (CUBE 640-40C, Coherent Inc.) lasers delivered through the rear port, with laser power controlled using an acousto-optic tunable filter (AOTFnC-400.650-TN, AA Opto-Electronic). Illumination was directed via a four-band excitation dichroic mirror (Di01-R405/488/561/635; Semrock), and emission signals were collected through wavelength-specific filters (FF01-525/50 for green and FF01-605/64 for red; Semrock) mounted on a motorized filter wheel (FW-1000, Applied Scientific Instrumentation) to enable rapid channel switching.

Two-color simultaneous imaging of *Arc* and *Actb* mRNAs was performed using a custom built wide-field fluorescence microscope constructed on an Olympus IX-81 platform equipped with two precisely aligned iXon Ultra EMCCD cameras (Andor), mounted on a multi-axis alignement stage. Channel-specific emission filters (FF03-525/50-25 for green and FF01-607/70-25 for red; Semrock) were positioned in front of each camera. Camera exposures were synchronized using TTL triggers generated by a DAQ board (Measurement Computing). The microscope was equipped with a piezoelectric stage (ASI) for rapid z-positioning and a Delta-T environmental chamber (Bioptech) for live-cell imaging. Two-color live imaging was performed in a single focal plane with frame acquisition at 10Hz.

### Glutamate Uncaging

Photolytic uncaging of MNI-caged-L-glutamate (Tocris) was used to locally stimulate NMDA receptors (NMDARs) at dendritic spines, as previously described^16^. Glutamate uncaging was performed along 5–10 μm distal dendritic segments using three sequential stimulation trains (separated by 2-minute intervals) consisting of low-frequency pulses (0.5 Hz) from a diffraction-limited 405 nm laser delivered at six sites positioned 0.5–1 μm from the dendritic shaft. This stimulation paradigm reliably induces NMDAR-dependent Ca²⁺ influx and triggers sustained structural long-term potentiation (sLTP) at stimulated spines.

During imaging, hippocampal neurons were maintained in Hibernate A Low Fluorescence Mg²⁺-free medium (BrainBits) supplemented with 2 mM MNI-caged-L-glutamate and 1.5–2 μM tetrodotoxin (TTX) (Tocris), with the stage-top incubator (Tokai Hit) set to 35 °C. Each uncaging sequence lasted approximately 30 seconds, after which 11 z-sections (400 nm step size) were acquired every 15 seconds over a total imaging period of 12-15 min. Z-stacks from each time point were processed using maximum-intensity projection in Fiji/ImageJ (NIH) and assembled into time-lapse movies for downstream analysis.

### Image Analysis (Fixed images)

#### Colocalization Analysis of sm-FISH Images

Z-stack images of neurites containing *Arc* and *Actb* mRNAs were maximum-intensity projected and background-subtracted using the Subtract Background function in Fiji. Actin FISH spots in the neurites were filtered and fitted to a 3D Gaussian distribution using the Fiji RS-FISH plugin to determine the coordinates of the mRNA. The intensity and the width of the 3D Gaussian were thresholded to exclude non-specific and autofluorescent particles. Similarly, the Arc FISH spots were analyzed using RS-FISH. Within a single neurite, the *Arc* coordinates were cross-referenced against the *Actin* coordinates using the 3D Euclidean distance formula. For each matched image, puncta were analyzed across the full field and within four spatial bins defined along the x axis relative to soma distance: 0-40, 40-80, 80-120, and 120-160 µm. *Actb* and *Arc* puncta were considered co-associated when their two-dimensional Euclidean separation was less than or equal to 0.25 µm (250 nm). Within each image and bin, we quantified the number of *Actb* and *Arc* puncta, and the number of puncta in each channel having at least one neighbor from the other channel within threshold, and the total number of *Act*-*Arc* pairs within threshold. Co-association frequency was defined as the total number of associated *Actb* and *Arc* puncta divided by the total number of puncta in that image and bin. To evaluate enrichment above chance, we generated image-level null distributions by bootstrap resampling. For each image, *Actb* puncta and *Arc* puncta were resampled independently with replacement from the observed coordinate sets for 5,000 iterations while preserving the number of puncta in each channel. Co-association frequency was recalculated for each resampled dataset and spatial bin. For each bin, the group-level null distribution was defined as the iteration-wise mean of the per-image null co-association frequencies across eligible images. Observed mean co-association frequency was compared with this null distribution to calculate the null mean, null standard deviation, 95% null interval, z score, and an empirical one-sided enrichment p value computed as (1+nnull≥obs)/(N+1).

Because distal bins often contained very few puncta, we examined puncta counts per image in each spatial bin before finalizing summary analyses. To reduce instability caused by sparse denominators, image-bin observations containing fewer than five total puncta were excluded from bin-specific summary statistics and visualization.

### Distance Analysis

Dendritic segments were isolated from images of neurons, with the selection consistently beginning in the most proximal part of the neurite. sm-FISH spots in the neurites were analyzed via the RS-FISH plugin in Fiji and were filtered and fitted to a 3D Gaussian distribution to determine the coordinates of both mRNA types. The distance of the mRNA coordinates from the most proximal center point of the neurite was calculated using the 2D Euclidean distance formula. The mRNA was separated into 30 um bins progressing from the soma and normalized to the total neurite length within each bin.

### Image Analysis (Live imaging)

Movements of *Arc* and *Actb* mRNAs in dendrites were analyzed using Kymographs generated by Kymolyzer^112^. To identify regions of interest, timelapse images were maximum-intensity projected; only dendrites containing at least one actively translocating punctum for both mRNA species were selected for analysis. Dendrites were traced from the soma to an unambiguous distal termination along the dendritic midline, with an average of three dendrites analyzed per image. Only dendrites that had at least 3 mRNAs of each species and a mixed population of both moving and stationary mRNAs were included for analysis. Branch points were handled by removing the shared root between two images to preserve the distance from the soma but prevent reanalysis of the same mRNA. To avoid redundant analysis of the same mRNA puncta at branch points, the shared root was assigned to only one path to preserve distance-from-soma while preventing double-counting. Preference was given to paths containing mRNAs that translocated beyond the bifurcation. Kymographs were generated using a line width of 20 pixels (1.3 µm) to encompass both mRNA signals along the same centerline. Individual mRNA puncta were manually traced on the kymograph and cross-validated against the original timelapse series.

The resulting trajectories were compiled using Kymolyzer, using a velocity threshold of greater than 0.2 µm/second between consecutive frames to distinguish between moving and resting states. All puncta with a mean velocity of greater than 0.2 µm/sec were categorized as moving particles. The motile fraction (the ratio of moving to total puncta) was calculated for each dendrite. Statistical significance for differences in mRNA velocity, displacement, and motile fraction was determined using the Kolmogorov-Smirnov test.

The analysis for the uncaging experiments were performed as before^16^ by counting the number of mRNAs in the uncaged region, and their timing of arrival recorded after the first uncaging train. The efficiency of localization refers to the frequency of successful RNA localization events following glutamate uncaging in each experiment (3 - 4 neurons per experiment). A successful localization event refers to the recruitment of the RNA to the uncaged region, whereby the RNA localizes for a minimum of 3 frames. The residence time is the duration the mRNA resides in the uncaged region during the imaging session.

## Acknowledgements

We thank Chiso Nwokafor for animal maintenance and Melissa Lopez-Jones for technical assistance. We thank all members of the Jovanovic, Singer, and Das laboratories and the members of the Einstein Program in RNA biology group, particularly U. Thomas Meier, and Robert Coleman for their helpful feedback. The work is supported by National Institutes of Health grant R01 NS083085 to R.H.S. and S.D.; R35GM152258 to M.J., K99NS135103 to E.D.M., T32GM007491 and T32AG023475 (R.C.) and F31NS137668 to J.R.

## Data and materials availability

The RNA-seq data generated in this study have been deposited in the Gene Expression Omnibus (GEO) under accession number GSE326430. Code used for data processing and analysis is publicly available on GitHub at github.com/cutleraging/Rogow-Doron-Mandel-Cutler-2026.

