## Supplementary Figures for "An integrated RNA-centric imaging and omics approach reveals distinct properties and composition of neuronal RNA granules"

A

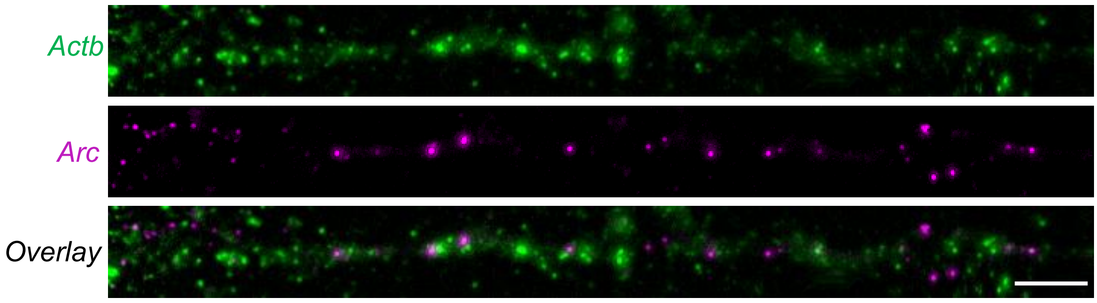


B

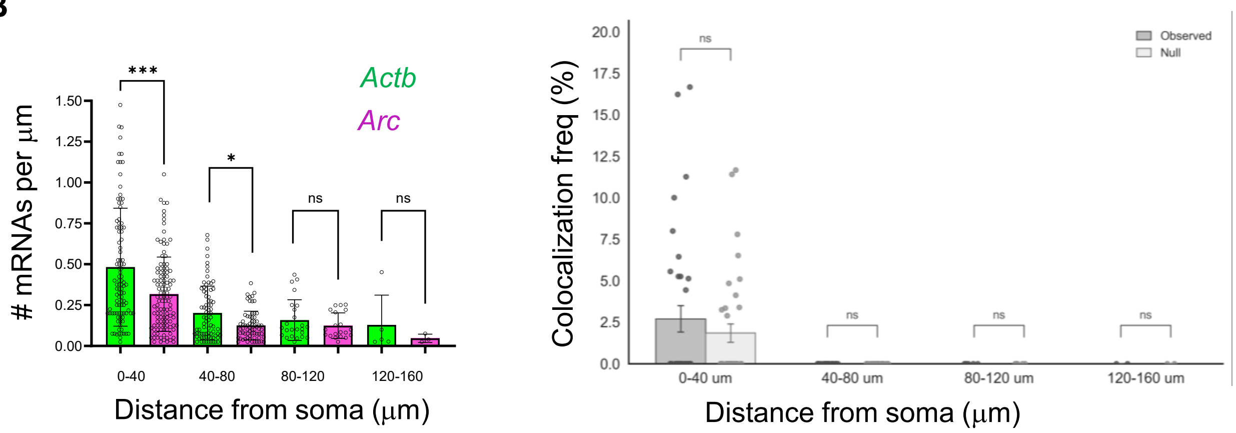


**Supplementary Figure 1. *Arc* and *Actb* mRNA association in fixed hippocampal neurons. (A)** Single molecule FISH for *Arc* and *Actb* mRNAs was performed one hour after stimulation. Representative images distinct *Arc* and *Actb* granules in dendrites. **(B)** RNA density as a function of distance from the soma (n = number of dendritic segments; 103 (*Actb*), 115 (*Arc*) for 0-40 bin, 79 (*Actb*), 67 (*Arc*) for 40-80 bin, 24 (*Actb*), 20 (*Arc*) for 80-120 bin, 5 (*Actb*), 3 (*Arc*) for 120-160 bin, Mann-Whitney test. **(C)** Frequency of co-localization of *Arc* and *Actb* mRNAs, Statistics: empirical one-sided resampling test based on a bootstrap null distribution. n= 35 dendrites (0 - 40 bin), 21 dendrites (40-80 bin), 5 (80 -120 bin) ***:p<0.001, *:p<0.05

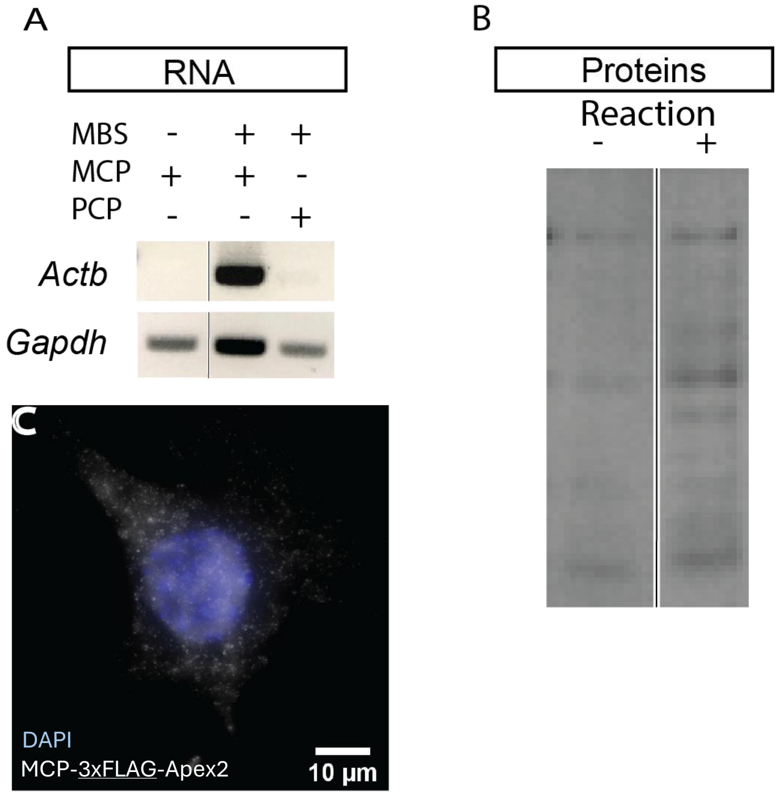


**Supplementary Figure 2. Validation of RNA and protein biotinylation of *Actb* mRNP using APEX in Mouse embryonic Fibroblasts (A)** APEX reaction biotinylates RNA in the *Actb* mRNP in mouse embryonic fibroblasts expressing integrated MCP-APEX2 lentivirus. Reverse Transcriptase PCR detection of *Actb* mRNA after streptavidin bead pulldown occurs only in samples with stem loops and MCP fused APEX2 (lane 2), but not when either the stem loops or MCP is missing (lanes 1 and 3). **(B)** APEX reaction yields more biotinylated proteins by whole cell lysate western blot with streptavidin-HRP. Lanes before and after reaction with APEX2. **(C)** MCP-3xFLAG-APEX2 is detected by immunofluorescence via anti-flag antibody when the construct is expressed in fibroblasts (White = APEX; Blue = DAPI).

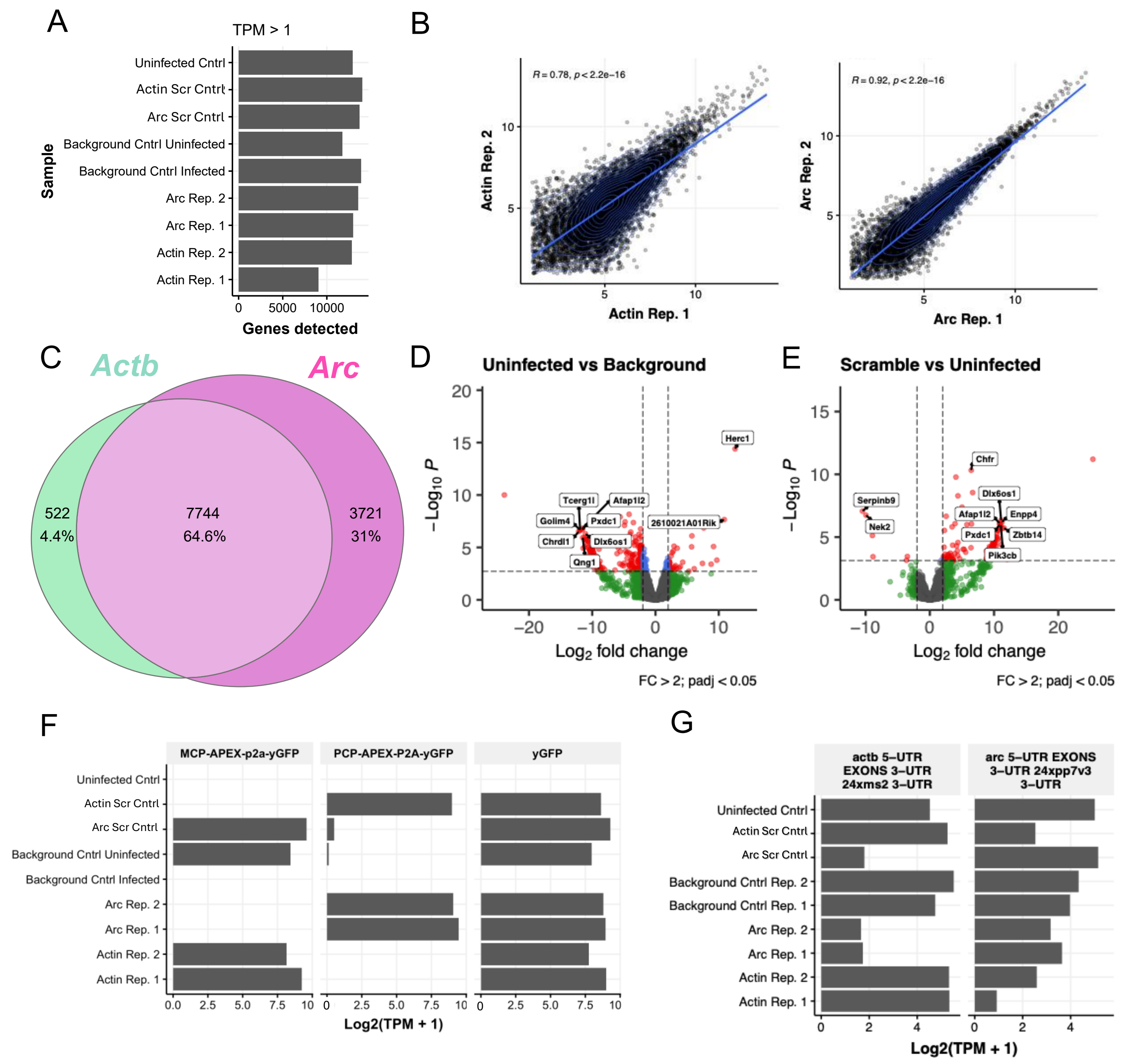


**Supplementary Figure 3. APEX-Seq quality control, related to Fig. 3. (A)** Number of genes detected with TPM>1. **(B)** Correlation of gene expression levels between biological replicates for *Actb* and *Arc* RNA proximity biotinylation. Statistics: Significance of Pearson R correlation coefficients for each batch using a two-sided t-test. **(C)** Overlap between genes detected in *Actb* and *Arc* samples following gene level filtering (Methods). **(D-E)** Volcano plots showing (D) the effect of the streptavidin bead pull down (uninfected vs. background) or (E) the effect of the MCP/PCP virus infection (scramble vs. uninfected). The significantly upregulated genes from both of these comparisons were used to filter false-positive hits from *Actb/Arc* enrichments. Red points are $Log_2$ fold change>2 and adjusted p-value<0.05. Statistics: p-values estimated using a Wald test followed by Benjamini-Hochberg correction for multiple hypothesis testing. **(F)** Expression levels of viral sequences specific and non-specific to each condition. **(G)** Expression levels of endogenous genes including stem-loop sequences

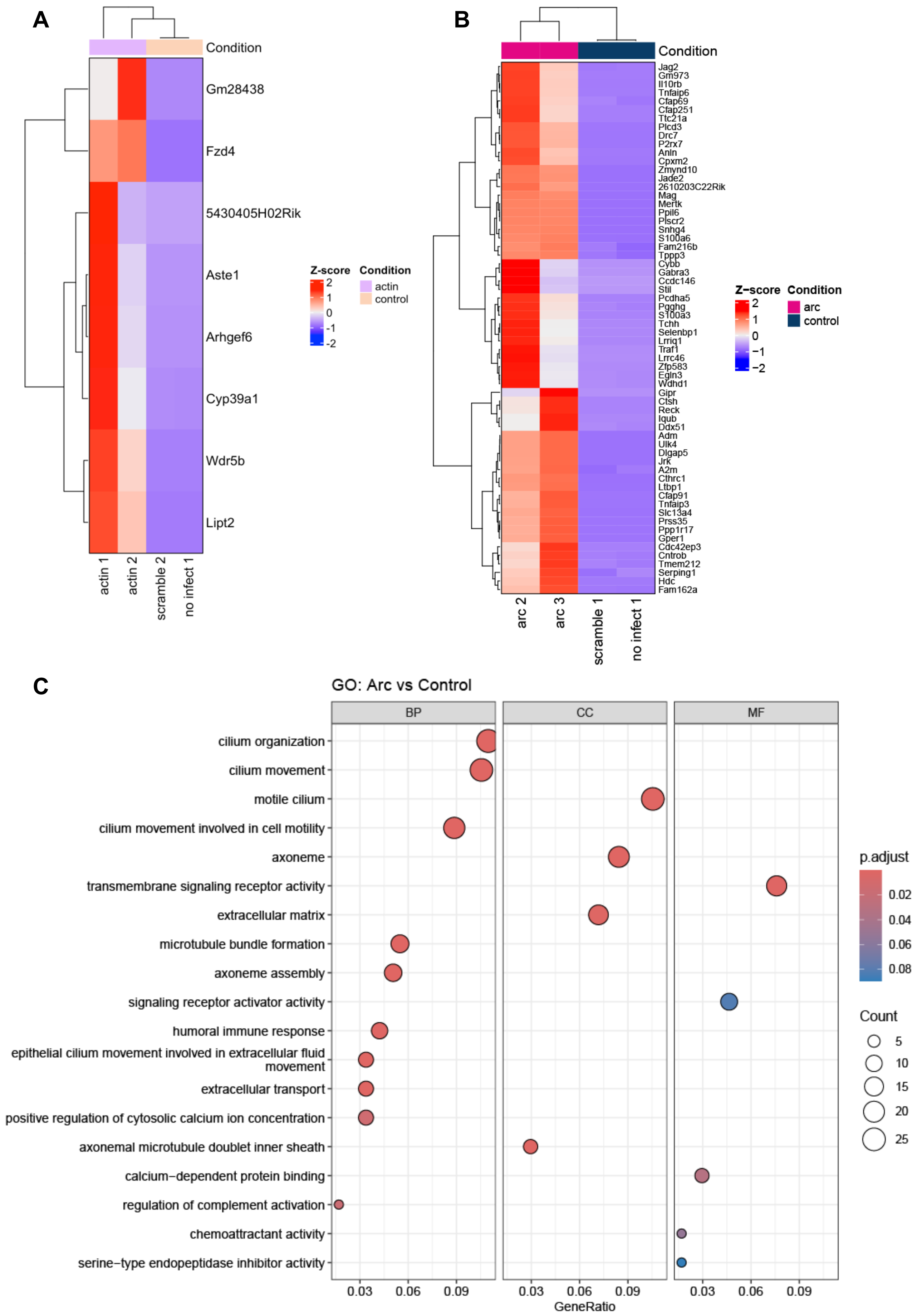


**Supplementary Figure 4. RNA Proximity Biotinylation enrichment, related to Figure 3. (A)** Heatmap and hierarchal clustering of significantly enriched genes ($Log_2$ fold change>2 and adjusted p-value<0.05) associated with *Actb* RNA. **(B)** Same as in A, but for *Arc*. **(C)** Over-representation analysis showing top 10 gene ontology enrichments for significantly enriched genes associated with *Arc* RNA. Statistics: adjusted p-values values were computed using a hypergeometric test followed by Benjamini–Hochberg false-discovery-rate correction.

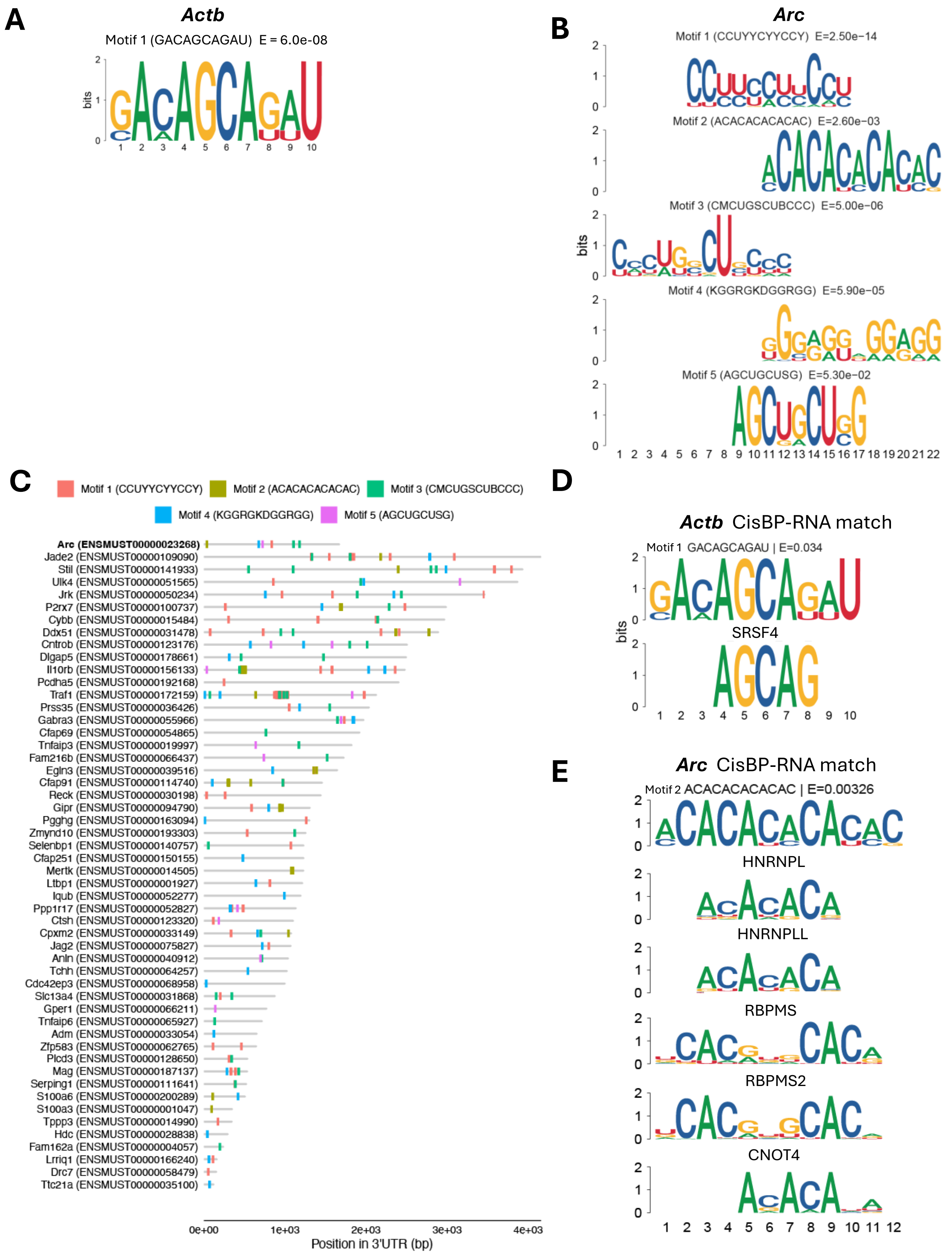


**Supplementary Figure 5. 3'UTR RNA Motif Enrichment, related to figure 3. (A)** *Actb de novo* motif enrichment results. Statistics: E-values were estimated by taking the motif's enrichment p-value and multiplying it by the number of motif tests performed, giving the expected count of equally strong motifs arising by chance. **(B)** *Arc de novo* motif enrichment results. Motif are aligned. Statistics: Same as in (A). **(C)** Full list of *de novo* motif occurrences in *Arc* and 3' UTR sequences for genes enriched from the *Arc* proximity biotinylation, IUPAC nomenclature (Fig. 3C). Motifs are shown with matches to 3' UTR region with q-value<0.1. Statistics: p-values are derived from the probability of equal or higher scoring matches under a background model, with multiple testing correction performed using Benjamini–Hochberg FDR. **(D)** Alignment of *de novo* motif 1 found in the 3'UTRs of significantly enriched *Actb* associated mRNAs and the RBP SRSF4. Statistics: E-value was estimated by taking the motif's enrichment p-value and multiplying it by the number of motif tests performed, giving the expected count of equally strong motifs arising by chance. **(E)** Alignment of *de novo* motif 2 found in the 3'UTRs of significantly enriched *Arc* associated mRNAs and the RBPs HNRNPL, HNRNPLL, RBPMS, RBPMS2, and CNOT4. Statistics: Same as in (D).

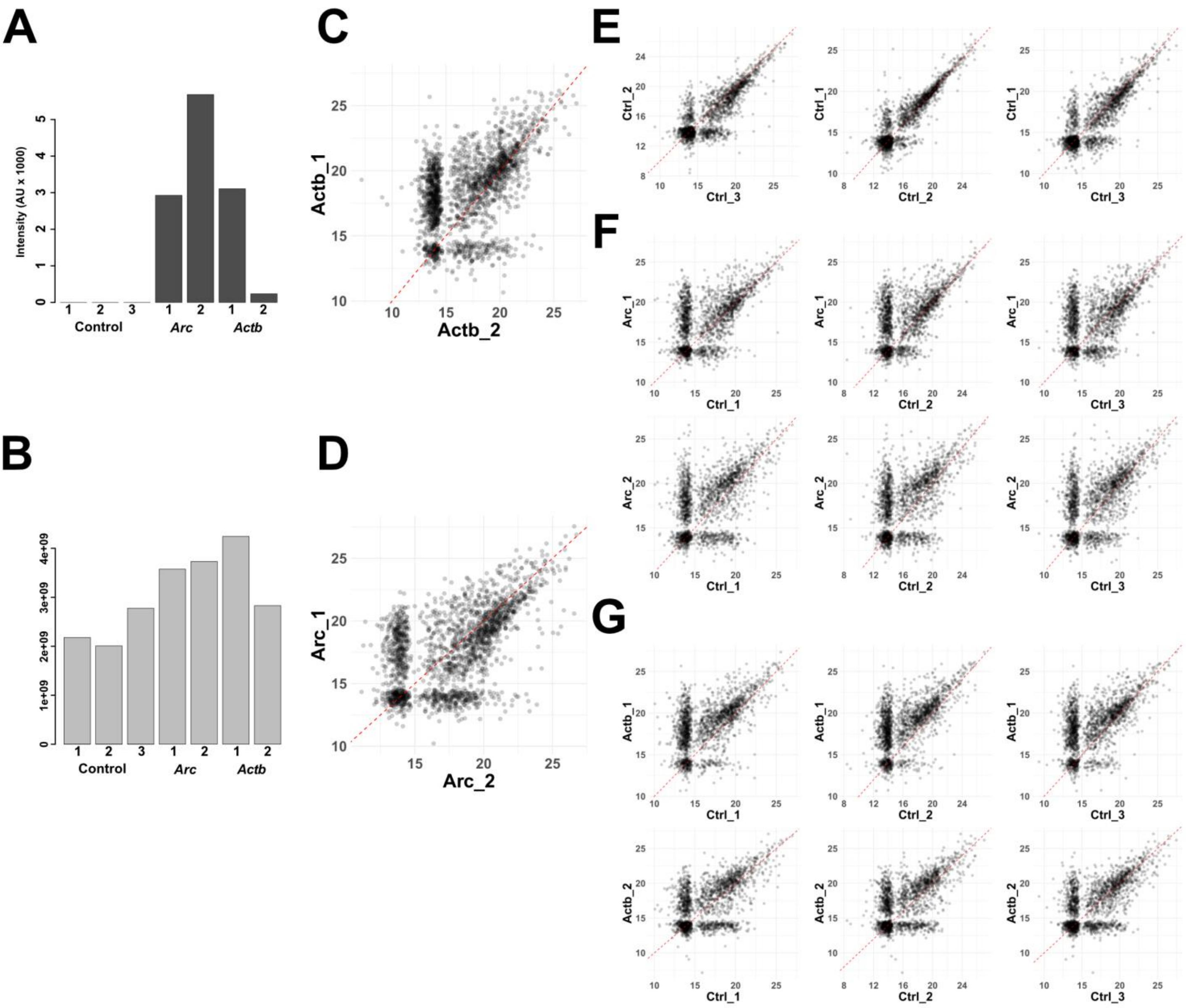


**Supplementary Figure 6. Protein proximity biotinylation - quality controls, related to Fig 4 (A)** Intensity of the transfected coat-protein-APEX2 fusion protein (based on common sequence between MCP-APEX2 and PCP-APEX2) in the enriched biotinylated proteomes across samples. **(B)** Total intensity measured (sum of all proteins per sample). **(C)** Scatterplot comparing the log2(Intensity) of the experimental *Actb* samples. **(D)** Scatterplot comparing the log2(Intensity) of the experimental *Arc* samples. (E) Scatterplot comparing the log2(Intensity) of the 3 negative control samples. **(F)** Scatterplot comparing the log2(Intensity) of each *Arc* experimental sample to each of the controls. **(G)** Same as (F), for *Actb* experimental samples.

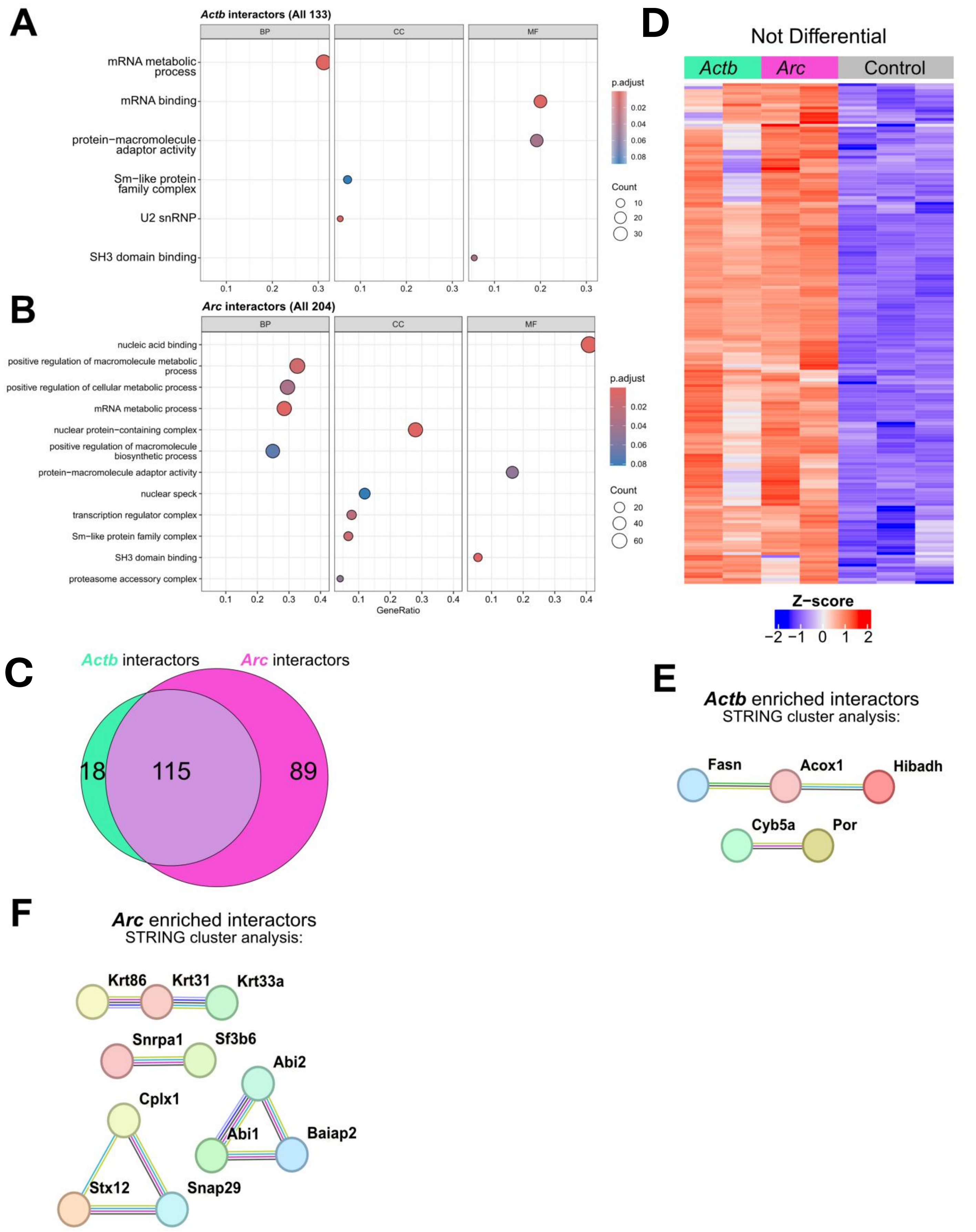


**Supplementary Figure 7. Protein proximity biotinylation – extended analysis, related to Fig 4 (A)** GO term enrichment based on the 133 proteins identified as *Actb* interactors. **(B)** GO term enrichment based on the 204 proteins identified as *Arc* interactors. **(C)** Euler plot showing the overlap in identified interactors between *Arc* and *Actb* vs control. **(D)** Heatmap representation of the signal of *Actb* or *Arc* interactors that were not significantly differential between *Arc* and *Actb* samples in the direct (*Arc vs Actb)* comparison. **(E)** STRING analysis for the group of *Actb* enriched interactors from the direct comparison (showed in main Figure 4D-E). Clusters based on interactions between genes. Interaction lines: Purple = experimentally determined, green = text mining, black = co-expression, blue = curated database, red = gene fusion. **(F)** STRING analysis for the group of *Arc* enriched interactors from the direct comparison (showed in main Figure 4D-E).
